# Revealing 3D cancer tissue structures using holotomography and virtual hematoxylin and eosin staining via deep learning

**DOI:** 10.1101/2023.12.04.569853

**Authors:** Juyeon Park, Su-Jin Shin, Minji Kim, Geon Kim, Hyungjoo Cho, Dongmin Ryu, Daewoong Ahn, Ji Eun Heo, Inyeop Jang, Hyun-seok Min, Kwang Suk Lee, Tae Hyun Hwang, YongKeun Park

## Abstract

In standard histopathology, hematoxylin and eosin (H&E) staining stands as a pivotal tool for cancer tissue analysis. However, this method is limited to two-dimensional (2D) analysis or requires labor-intensive preparation for three-dimensional (3D) inspection of cancer tissues. In this study, we present a method for 3D virtual H&E staining of label-free cancer tissues, employing holotomography and deep learning. Holotomography is used to measure the 3D refractive index (RI) distribution of the label-free cancer slides. A deep learning-based image-to-image translation framework is integrated into the resulting 3D RI distribution, enabling virtual H&E staining in 3D. Our method has been applied to colon cancer tissue slides with thicknesses up to 20 μm, with conventional chemical H&E staining providing a direct validation for the method. This framework not only bypasses the conventional staining process but also provides 3D structures of glands, lumens, and individual nuclei. The results demonstrate enhancement in histopathological efficiency and the extension of the standard histopathology into the 3D realm. To validate the repeatability and scalability of the approach, we applied the framework to the gastric cancer slides obtained from different institute and imaging devices.

## Introduction

Histopathology is a procedure involving the microscopic examination of tissue samples to study the structural cellular alterations in diseased tissues. Standard histopathological procedures meticulously stain tissue slides with hematoxylin and eosin (H&E), a widely used technique for enhancing the visibility of the cellular and tissue structures when examined under bright-field (BF) microscopes. The standard procedures, employed for centuries, provide valuable insights into microscopic structural changes in tissues, facilitating diagnosis, grading, and classification of diseases (Fig. 1a).

**Fig.1.**
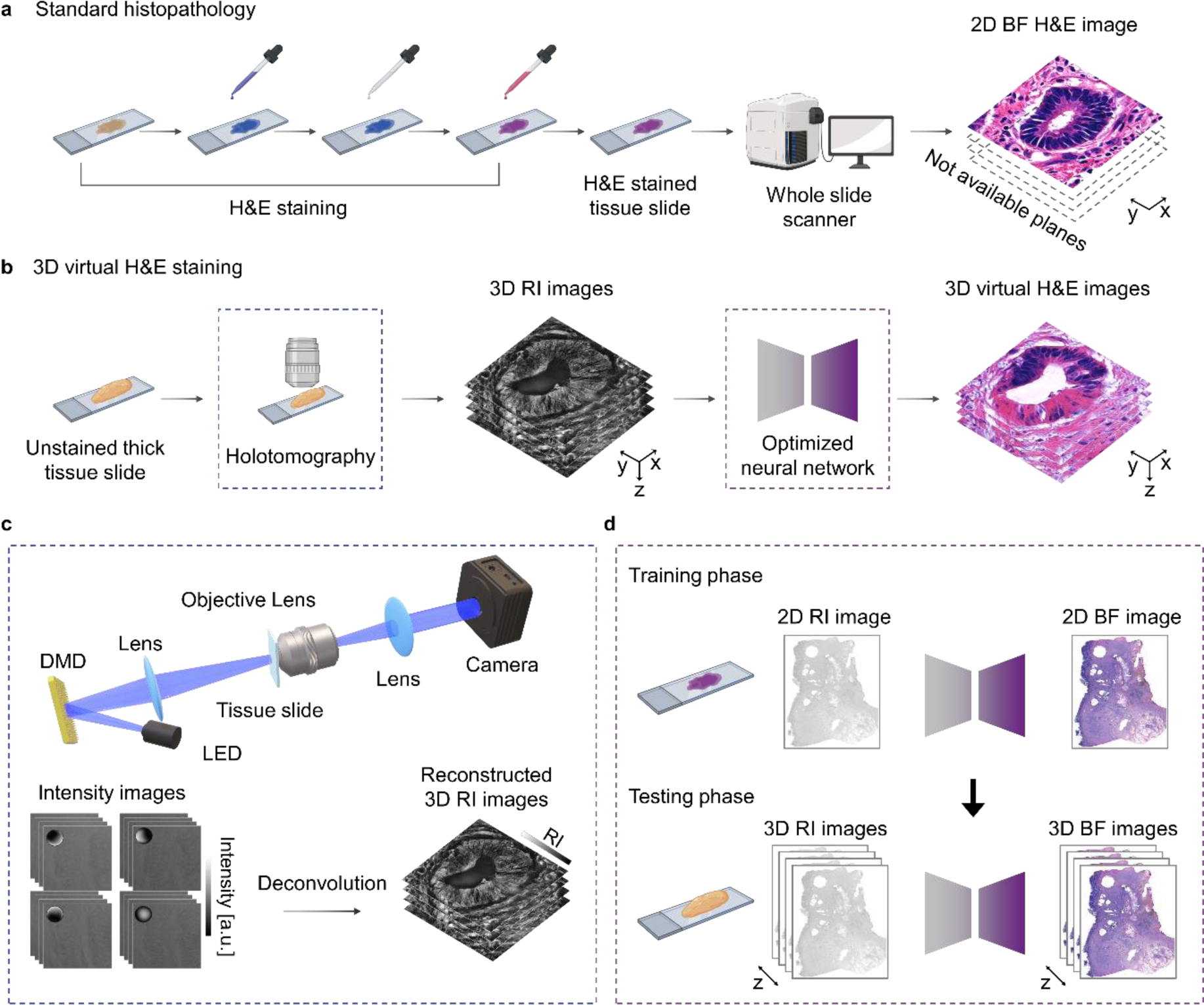
Overview of the proposed framework. **a.** Procedures for standard histopathology including H&E staining and imaging under BF microscopy. **b**. The proposed framework of 3D virtual H&E staining. Integration of holotomography and deep learning enabled label-free 3D virtual H&E staining of thick colon cancer tissue slides. **c**. Optical setup for holotomography and reconstruction procedures. Raw intensity images are deconvolved with the optical transfer functions, reconstructing 3D RI images. **d**. 3D Virtual H&E staining framework. The neural network is trained to map between RI and BF images using the conventional thin, H&E-stained colon and tissue slides during the training phase. The trained neural network is directly applied to the 3D RI images obtained from label-free, 10 and 20 μm-thick colon cancer tissue slides.

However, the standard histopathology procedure is limited to two-dimensional (2D) analysis. Although biological tissues inherently possess three-dimensional (3D) structures, standard histopathology has limitations in providing the 3D histopathological structures because of the challenges posed by BF microscopy and H&E staining when dealing with thick 3D tissue samples. Recent studies have highlighted the critical need for 3D histopathological analysis in histopathology, offering insights into 3D features such as vascular and neural structures^1^, tissue heterogeneity^2,3^, and cancer grading^4,5^. However, the current implementation of 3D histopathological analysis relies on heavy sample preparation steps, including hundreds of sectioning and H&E staining or optical clearing.

To mitigate the resource-intensive demands of 3D histopathology, numerous efforts have been made to eliminate staining procedures. Notably, quantitative phase imaging (QPI) techniques are one endeavor to investigate biomedical specimens in a label-free manner, which quantitatively measures the optical phase delay of the specimen without staining^6^. With the advantages of label-free imaging capabilities and quantitative measurements, QPI has become an invaluable tool for investigating histopathological specimens, including structures of diverse cancers^7–16^, Alzheimer’s disease^17^, and Crohn’s disease^18^. While the studies predominantly focused on 2D specimens, recent advancements in 3D QPI techniques are now being applied to various histopathological specimens^19–22^.

While using label-free imaging techniques, QPI images may lack subcellular specificity. To address this limitation, deep learning techniques have been actively integrated with QPI images to enhance subcellular specificity^23^. Among the various deep learning techniques for interpreting QPI images, the concept of virtual staining is gaining attention. Virtual staining involves an image-to-image translation framework, using deep learning to convert label-free images to the desired stained images. This approach has been applied to diverse histopathological slides and target stains. For instance, unsupervised deep learning has successfully generated immunohistochemistry-stained images from phase images of kidney tissue slides^24^. Additionally, supervised deep learning enabled virtual staining of nuclei and F-actin using the label-free phase images of kidney tissue slides^25^. Furthermore, Phasestain has generated H&E, John’s, and Masson’s trichrome stained images from label-free phase images of skin, kidney, and liver tissue, respectively^26^. These studies have demonstrated the potential of virtual staining to make standard histopathology more efficient by eliminating the staining procedures and replacing them with computational staining. However, it is important to note that these research efforts have primarily focused on 2D applications.

In this study, we aim to address two key challenges–confinement in two-dimension and resource-intensive H&E staining procedures–by employing holotomography and deep learning. Our approach involves the elimination of traditional H&E staining procedures and the extension of H&E stained images into three-dimensional, providing 3D H&E stained structures of colon cancer tissue slides without the need for staining (Fig. 1b). Holotomography, a 3D QPI technique, measures the label-free 3D refractive index (RI) distribution of the sample with high spatial resolution and optical sectioning capabilities. To realize the holotomography, we utilized the deconvolution phase microscopy which involves the deconvolution of intensity images using optical transfer functions to capture the 3D RI distribution of specimen without external staining^27^ (Fig. 1c). A deep learning technique for image-to-image translation is integrated with the holotomography to generate virtual H&E stained images from label-free RI images in 3D (Fig. 1d). The framework includes training and testing phase. During the training phase, the neural network learns to map between the RI images and H&E stained images obtained from a standard histopathological slide. Once the training is completed, the optimized neural network is directly applied to the 3D RI images acquired from 10 and 20 μm-thick, label-free colon cancer tissue slides, resulting in the 3D virtual H&E stained images.

## Results

### Network training

To train the network, we prepared the paired training dataset from a 4 μm-thick, H&E-stained colon cancer tissue slide (Supplementary Fig. 1a). We first measured the slide using holotomography to obtain RI images. It is important to note that the resulting 3D RI images were all-in-focused to ensure that the cellular details from slightly different axial positions could be integrated into a single focal plane^28^ (See Methods). Subsequently, the same slide was imaged using a whole slide scanner (WSS) to obtain the H&E-stained BF images. Image registration was performed between the BF images obtained from WSS and RI images measured using holotomography based on the spatial transform network^29^, and we cropped the images into 1,024 × 1,024-pixel patches, resulting in the paired dataset (See Methods). A total of 2,538 patches are prepared and the dataset is divided into 1,996 and 542 patches for training and validation, respectively.

For the architecture of the neural network, we employed a conditional generative adversarial network to train the model (Supplementary Fig. 1b). The network consists of a generator and discriminator, which are adversarially learning to map RI images to BF images. Scalable neural architecture search (SCNAS) has been used as the generator, an optimized architecture for 3D medical image datasets^30^ (Supplementary Fig. 2a). The discriminator consists of five convolutional layers (Supplementary Fig. 2b). While the generator creates the virtual H&E stained images from input RI images, the discriminator’s role is to distinguish the created image from the ground truth images when provided with input images.

To evaluate the performance of the trained networks, we compared virtual H&E images and ground truth images. First, we obtained a wide-field, all-in-focused RI image from a different region of the same slide used for training (Fig. 2a). The wide-field RI image revealed various histological features, including glands (Figs. 2a(ⅰ–ⅴ)) and stroma (Fig. 2a(ⅵ)). Subsequently, the wide-field RI image is cropped into 1024 × 1024-pixel patches with a 50% overlap and we fed the patches into the trained network. The generated H&E images are stitched back to reconstruct the wide-field image using the ImageJ plugin^31^ (Fig. 2b). To directly compare the virtual H&E images with ground truth images, we obtained the WSS image of the corresponding region (Fig. 2c). The results demonstrate that the trained network successfully predicted most of the histological and anatomical features including glands (Figs. 2b (ⅰ–ⅴ) and 2c (ⅰ–ⅴ)) and stroma (Figs. 2b (ⅵ) and 2c (ⅵ)). For the quantitative assessment, the structural similarity index measure (SSIM) values were calculated for each selected region of interest (See Methods).

**Fig. 2.**
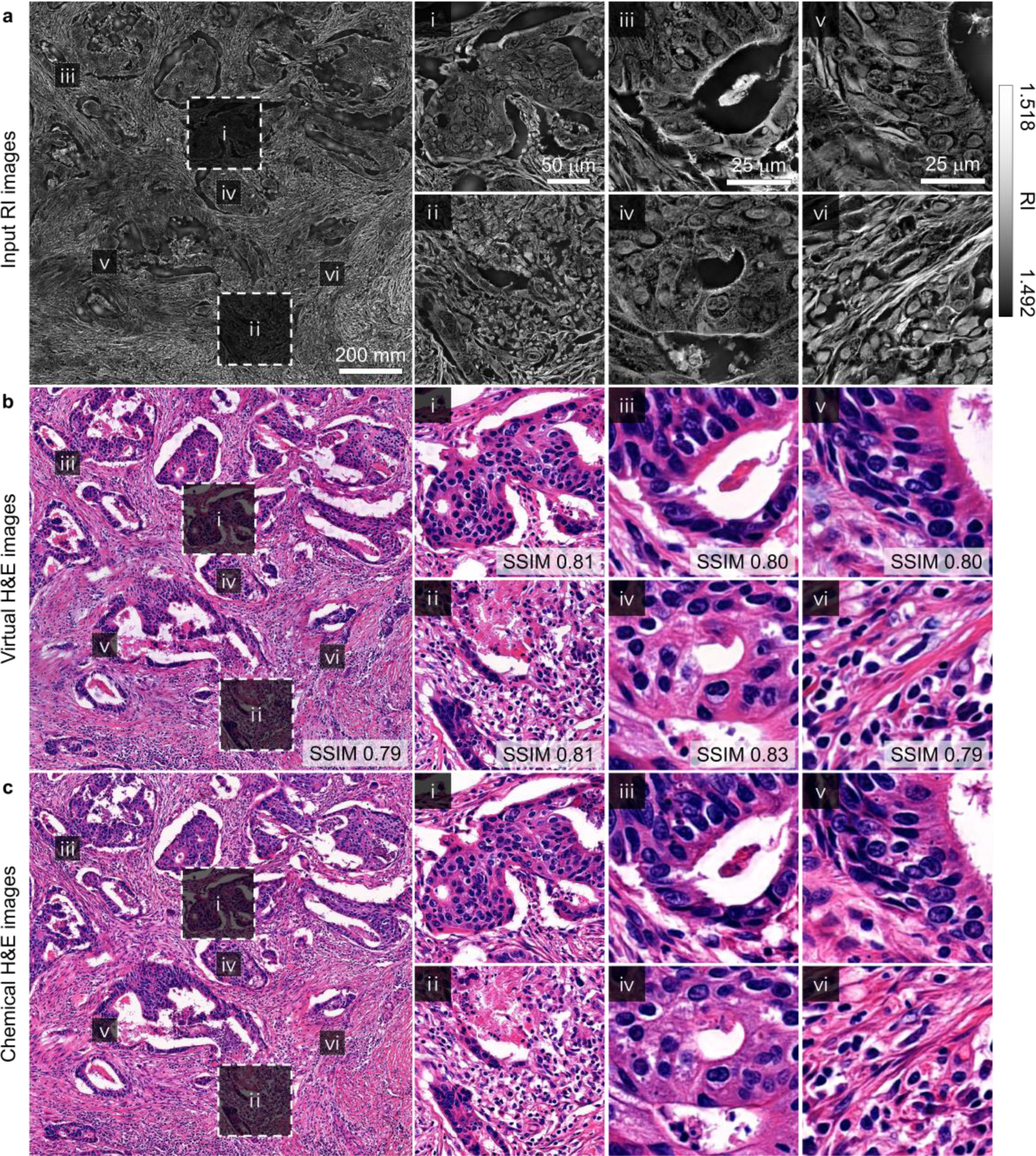
Validations of the trained network with a 4 μm-thick, H&E-stained colon cancer tissue slide. **a,** Wide-field RI image obtained from a 4 μm-thick, H&E-stained colon cancer slide and its detailed images. **b,** Wide-field H&E stained images generated by the trained neural network and its detailed images. **c,** Ground truth H&E images obtained using a WSS and its detailed images.

### 3D label-free RI distribution of thick colon tissue slides

To apply the trained neural network to label-free thick colon cancer tissue slides, we acquired 3D RI images from two colon cancer slides obtained from different patients. The slides were prepared with thicknesses of 10 and 20 μm, which are significantly thicker than the 4 μm-thick conventional histopathology slides, without any staining. We acquired approximately 1.2 × 1.2 mm regions of 3D RI images from each slide using holotomography (Fig 3). The wide-field RI images effectively depicted the anatomical structures, including glands and lumens, which are commonly found in colon cancer slides (Figs. 3a and 3c). Focusing more on the anatomical structures, we zoomed into the 3D structures of glands and lumens (Figs. 3b and 3d). Within these RI images, the lumen structures with vacant regions are distinctly clarified, as well as visualizing the surrounding glandular structures. Notably, the 3D structures exhibit the warping of lumens and surrounding glands along the axial axis. This dynamic presentation provides clear evidence of 3D reconstruction of the anatomical structures.

**Fig. 3.**
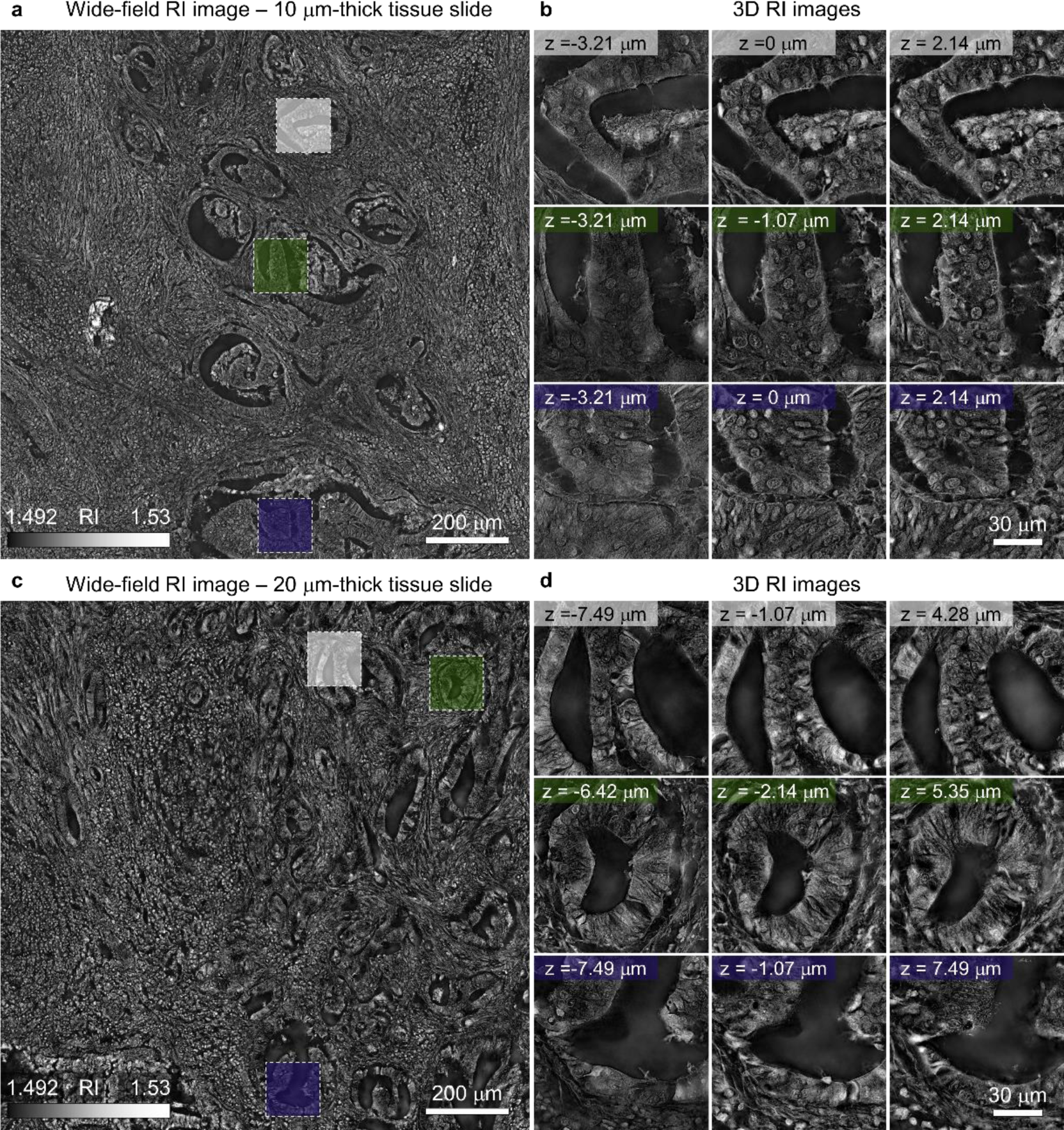
Label-free 3D RI images of thick colon cancer tissue slides. **a, c.** Wide-field RI images were obtained from the label-free 10 (a) and 20 μm-thick colon cancer tissue slides (c) using holotomography. A single section of 3D RI image is presented. **b, d.** Detailed 3D RI images of glands and lumens obtained from 10 (b) and 20 μm-thick colon cancer tissue slides (d).

### 3D virtual H&E images predicted by the trained network from label-free RI images

To generate the 3D H&E images from the label-free thick tissue slides, we employed the 3D RI images acquired from 10 μm and 20 μm-thick colon cancer tissue slides as input for our trained network. Specifically, we cropped each wide-field RI image from every axial position into 1024 × 1024-pixel patches with 50% overlap. These patches were used as input to the trained neural network, generating H&E images for each patch. The predicted H&E patches were stitched back to form a wide-field H&E images (Figs. 4a and 4c). By performing this prediction for all axial positions, we successfully generated 3D H&E images from the label-free thick colon cancer tissue slides.

**Fig. 4.**
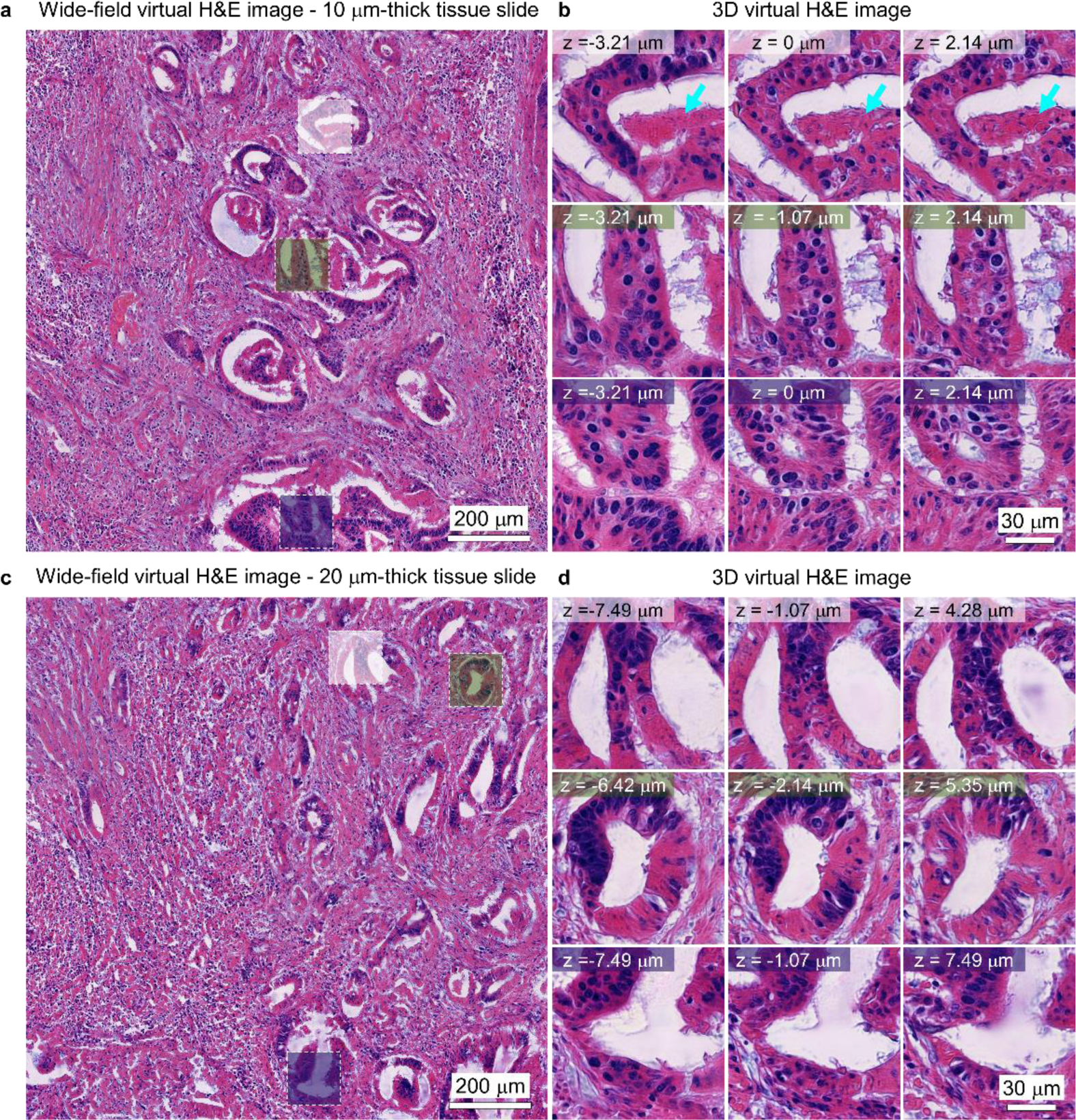
3D virtual H&E images predicted using trained neural network. **a, c.** Wide-field H&E images predicted from the label-free 10 (a) and 20 μm-thick colon cancer tissue slides (c). A single section of the 3D H&E image is presented. **b, d.** Detailed images of 3D H&E images predicted from 10 (b) and 20 μm-thick colon cancer tissue slides (d). Cyan arrows indicate the necrotic structures.

For both slides, anatomical structures including glands and lumens are successfully predicted as H&E stained images from the label-free RI images as shown in detailed images (Figs. 4b and 4d). Lumen structures with void areas as well as the surrounding glands are clearly identified. Additionally, the necrotic regions within the gland are clarified as indicated by cyan arrows. Building on the patterns observed in the RI images, the warping of the glands and lumens along the axial axis is consistently demonstrated in the predicted H&E images.

### Analyzing 3D histological features from the 3D virtual H&E images

To explore the detailed 3D outcomes of our trained neural network, we examined histological features in 3D (Fig. 5). In the 10 μm-thick sample, cross-sectional views revealed 3D gland and lumen structures (Fig. 5a). Void structures of lumens were continuously constructed along the *z*-axis, with width alterations in cross sections indicating the 3D construction of lumens. Notably, these cross sections also displayed the 3D structures of surrounding glands and the distribution of nuclei within the glands.

**Fig. 5.**
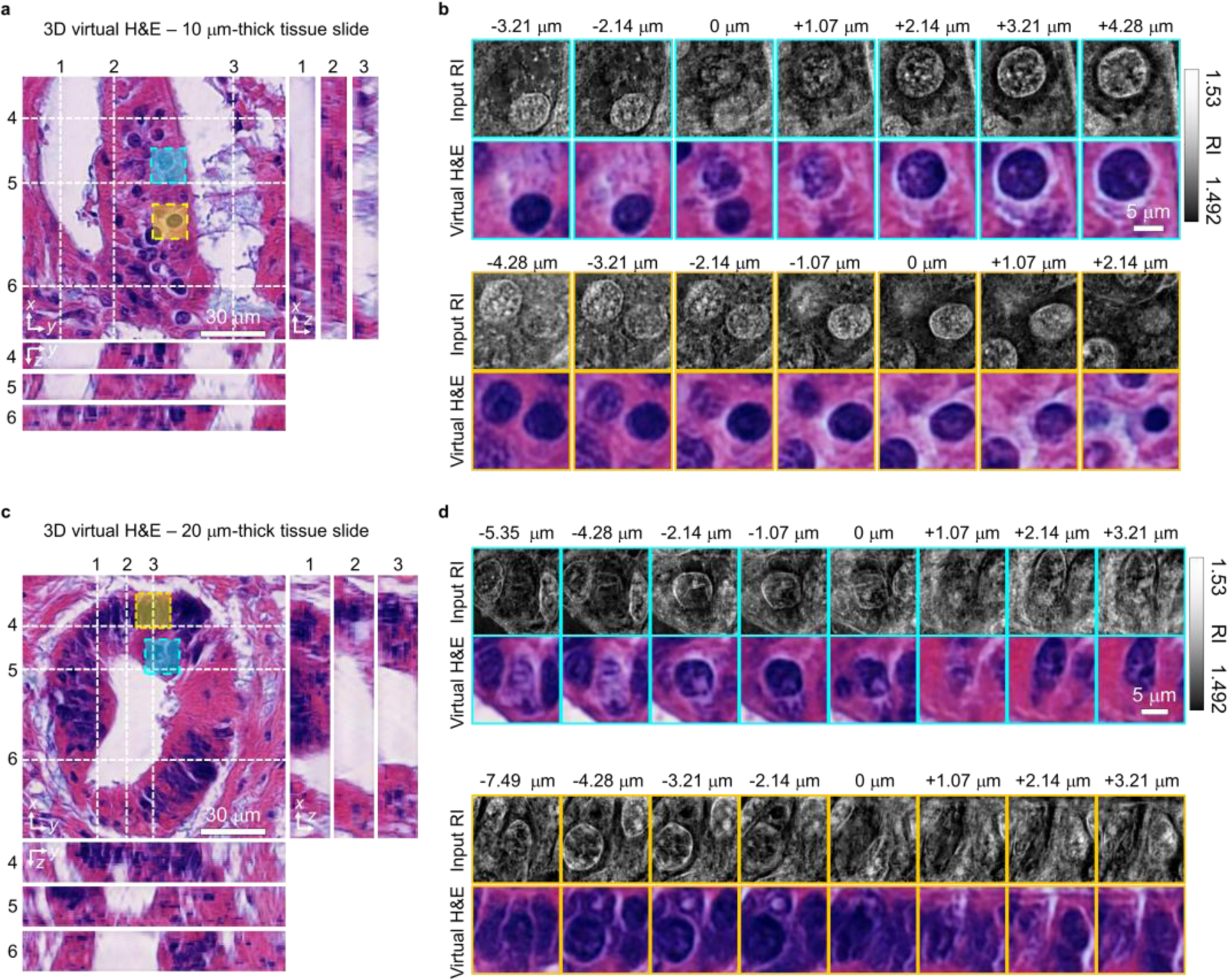
3D histological features of colon cancer tissue. **a, c.** The *x*-*y*, *y*-*z*, and *x*-*z* cross sections of predicted 3D H&E images of 10 (a) and 20 μm-thick colon cancer tissue slides (c). **b, d.** Detailed images to visualize individual nuclei in input RI images and corresponding predicted H&E images from 10 (b) and 20 μm-thick tissue slides (d).

Focusing on the scale of individual nuclei, we zoomed into 3D H&E images and compared them with input RI images (Fig. 5b). In our direct comparison, we observed circular structures of the nucleus in the input RI images, which is consistent with the predicted H&E images of blue circular morphologies. As the z-position changes, nuclei dynamically alter size and shape–contracting or expanding–indicating 3D construction of the nucleus. Furthermore, we examined the relative 3D distribution among nuclei by tracking the emergence and disappearance of some nuclei along the *z*-axis. This dynamic presentation provides a comprehensive understanding of the 3D arrangements of nuclei within the glands.

From the 20 μm-thick tissue, we similarly observed the 3D structures of glands and lumens (Fig. 5c). The cross sections depicted the 3D architecture of void structures of the lumen, exhibiting structural variations of widening, shrinking, or warping along the z-axis. Additionally, the cross sections of the surrounding glands revealed the 3D structures of the glands, along with the 3D distribution of nuclei within the glandular structures.

A detailed comparison between the input RI images and the predicted 3D H&E images provided 3D analyses of individual nuclei (Fig 5d). Specifically, as the axial position varied, some nuclei became visible, others disappeared, and certain nuclei underwent changes in both shape and size. This dynamic presentation unveils both 3D relative distribution among nuclei and 3D structures of individual nuclei within the glands.

### Validation of the virtual H&E images with the chemically stained slides

We further validated the capabilities of our method in the 3D histopathology of thick colon cancer tissue slides. To achieve this, we directly compared the 3D virtual H&E images to the chemically stained images of thick tissue slides (Fig. 6). After measuring label-free RI images of the two thick colon cancer slides, we chemically stained the same slide and obtained images using WSS. Because the WSS images are only available in 2D, we applied minimum intensity projection to the 3D virtual H&E images along the z-axis to ensure a fair comparison.

**Fig. 6.**
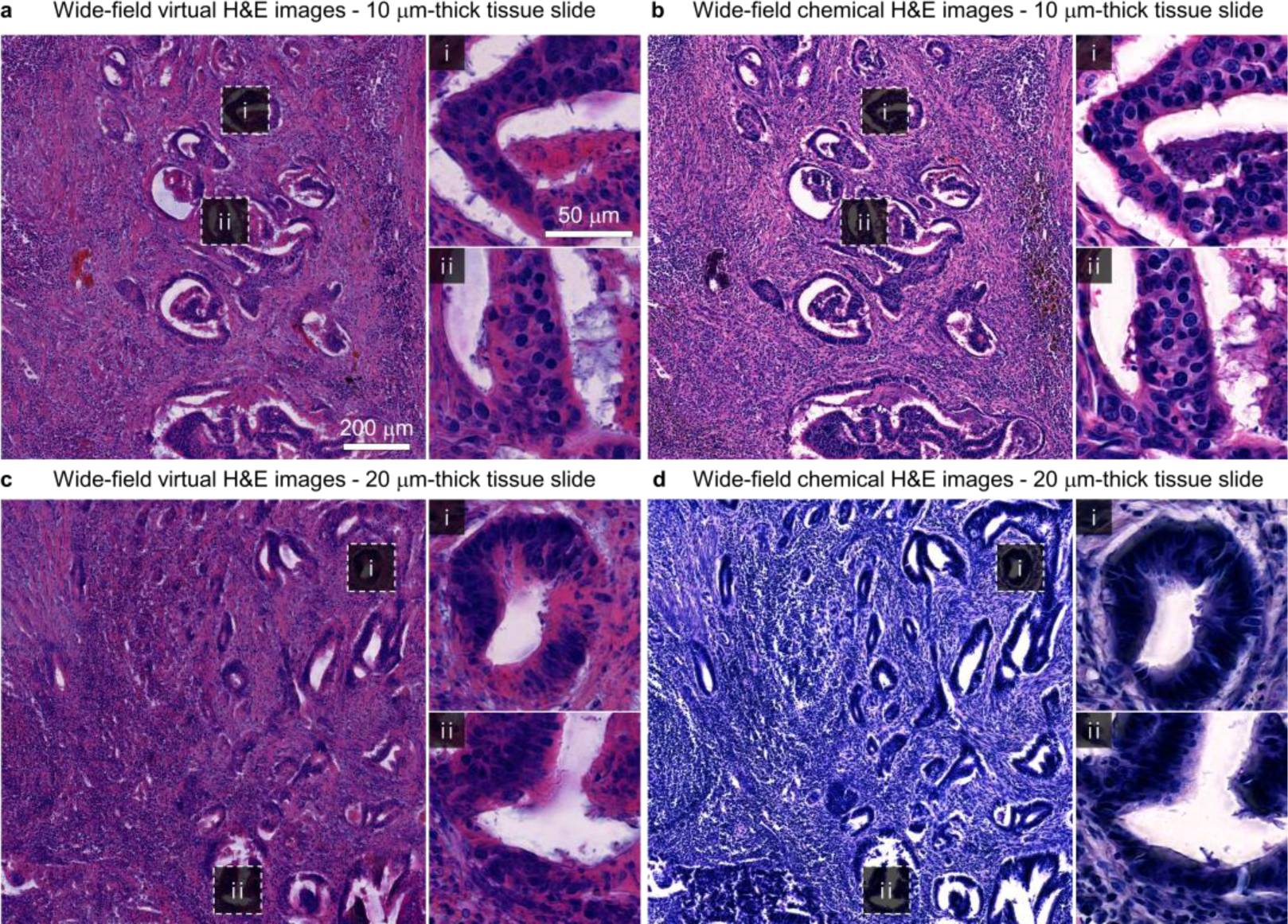
Validations using the standard histopathology procedures. **a, c,** Minimum intensity projection of predicted 3D H&E images from label-free 10 (a) and 20 μm-thick colon cancer tissue slide (c). **b, d,** Same 10 (b) and 20 μm-thick tissue slides (d) were stained with H&E and imaged using WSS. Detailed images of selected glandular structures are presented (ⅰ, ⅱ).

The comparison between virtual and chemical H&E images for the 10 μm-thick tissue slide highlighted the reliability and fidelity of the proposed virtual staining framework. The analysis of the two wide-field images exhibited the accurate reproduction of anatomical features by the trained network (Figs. 6a and 6b). The anatomical structures including glands and stroma are visible in the virtual H&E stained images. To provide a detailed examination, we zoomed the glandular structures, revealing comparable nucleus distributions to the chemically stained slides (Figs. 6a (ⅰ, ⅱ) and 6b (ⅰ, ⅱ)).

Comparing virtual and chemical H&E for the 20 μm-thick tissue slides demonstrated the robustness of the virtual staining framework, even for sample thickness where the chemical staining becomes abnormal. The colorimetric characteristics of virtual H&E remain consistent with minimal deviation, while chemical staining resulted in a distinctly different color distribution. Nevertheless, we could compare the anatomical features between virtual and chemical H&E images (Figs. 6c and 6d). The structures of glands and stroma were successfully reproduced by the trained network. For a detailed investigation of glandular structures, we zoomed into the selected glands, clearly illustrating the consistency of lumens and surrounding glandular structures (Figs. 6c (ⅰ, ⅱ) and 6d (ⅰ, ⅱ)). All regions indicated with arrows and boxes are confirmed by the experienced pathologist (SJS).

### Inter-organ, inter-institutional, and inter-device validation of 3D virtual H&E staining

To ensure the scalability and repeatability of our framework, it is crucial to demonstrate the direct application of our virtual staining technique across diverse sample origins, institutional settings, and imaging devices. To achieve this, we conducted virtual staining procedures on gastric cancer slides obtained from the Mayo Clinic. The procedures for generating the training dataset remained consistent with those used for the colon cancer dataset, differing only in the number of slides and imaging devices used (See Methods). This resulted in a dataset consisting of 9,231 patches, with 7,002 patches allocated for training and 2,231 patches for validation. Following training, SSIM values were computed for each selected window to quantitatively evaluate the network’s performance (Supplementary Fig. 3). Additionally, we utilized identical architecture and hyperparameters of the neural network as those employed for training on colon cancer data, ensuring consistency and robustness of our neural network architecture.

Following training, we prepared a 20 μm-thick gastric cancer tissue slide without any staining. Then, we acquired 3D RI images spanning approximately 1.92 × 1.22 mm regions using holotomography (Fig. 7a). These wide-field RI images clearly depicted anatomical structures such as smooth muscle and blood vessels, with vascular structures displaying individual cells within circular surrounding tissue structures, alongside wavy textures in the smooth muscle (Fig. 7b).

**Fig. 7.**
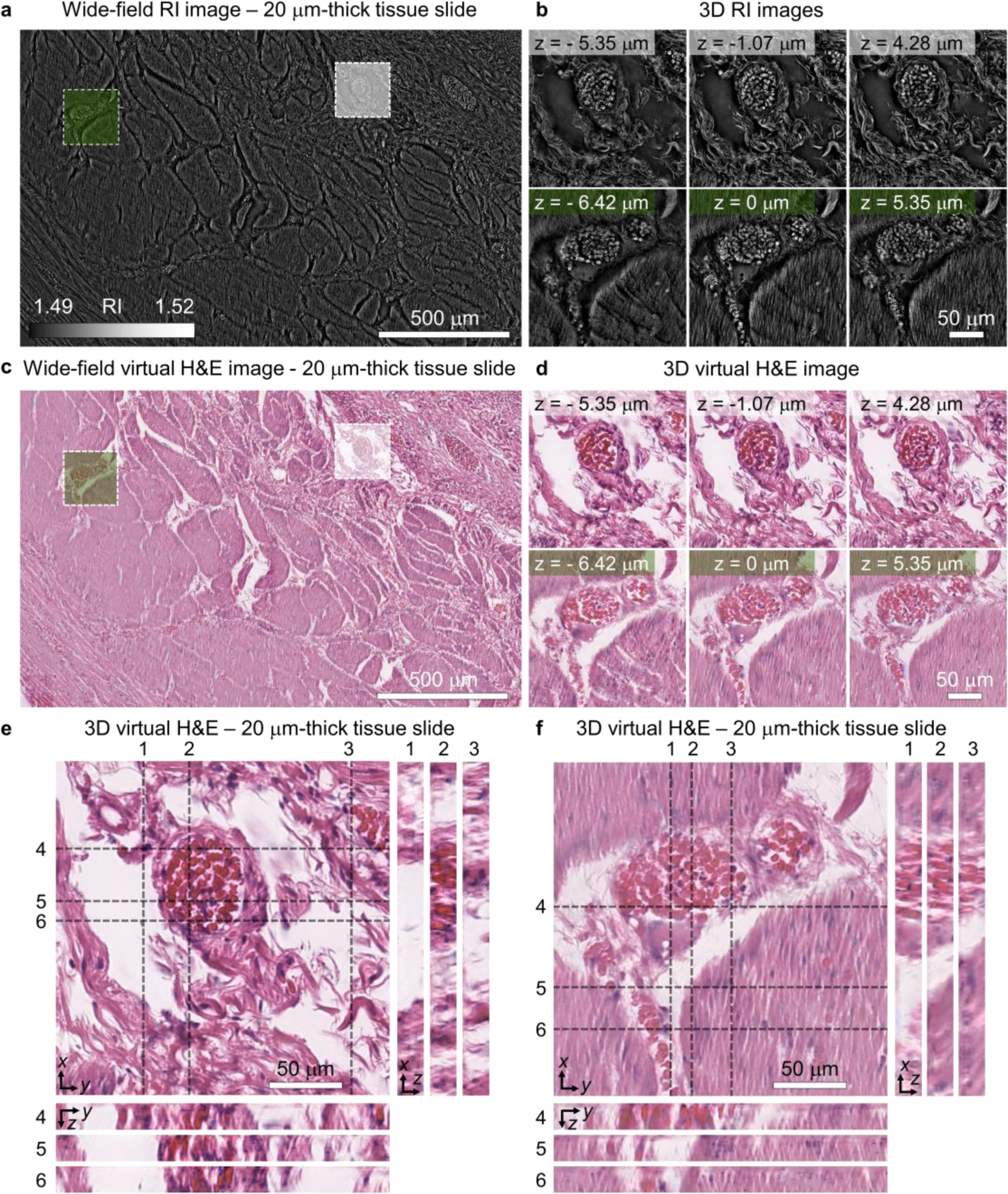
Validations of the virtual staining across different organ type, institute, and imaging device. **a.** A wide-field RI image was obtained from the label-free 20 μm-thick gastric cancer tissue slide using holotomography. A single section of 3D RI image is presented. **b.** Detailed 3D RI images of muscular and vascular structures obtained from the 20 μm-thick gastric cancer tissue slide. **c.** A wide-field H&E image predicted from the label-free 20 μm-thick gastric cancer tissue slide. A single section of the 3D H&E image is presented. **d.** Detailed images of 3D H&E images predicted from the 20 μm-thick gastric cancer tissue slide. **e, f.** The *x*-*y*, *y*-*z*, and *x*-*z* cross sections of predicted 3D H&E images of the 20 μm-thick gastric cancer tissue slide.

Applying the same cropping and stitching procedures used for colon cancer data, we inputted the 3D RI images into our trained network to generate virtually stained 3D H&E images of the gastric cancer slides (Fig. 7c). Notably, the network effectively predicted most anatomical structures, including smooth muscle and vascular structures, as evidenced in wide-field images. Detailed examination revealed a successful depiction of vascular structures as red blood cells with surrounding circular connective tissue structures and wavy textures of smooth muscle (Fig. 7d). Leveraging the 3D nature of the data, we observed changes in the cross-sectional morphology of vascular structures along the axial axis, as well as alterations in the distribution of red blood cells within it. Moreover, variations in the wavy texture of smooth muscle along the axial axis were visualized, highlighting the ability of virtual 3D H&E staining to reveal intricate 3D tissue textures.

To further elucidate structural features in 3D, we visualized cross sections of detailed images depicting vascular and muscular structures (Figs. 7e and 7f). Notably, these cross sections delineated continuous structural changes of vessels and muscles, exhibiting curved propagation along the axial axis. Additionally, void structures are clearly distinguished from the tissue regions, enhancing the clarity of tissue structures in three dimensions.

## Discussion

The proposed virtual H&E staining framework effectively addresses two limitations–labor-intensive sample staining procedures and 2D limited analysis–simultaneously. By utilizing a single routinely generated histopathological slide, we trained the neural network that translates RI images into H&E stained images. This trained network, when applied to 3D RI images of label-free thick tissue, successfully generated 3D H&E images. This approach not only eliminates the need for staining procedures and significantly reduces the number of slicing processes but also provides comprehensive 3D glandular structures for thick colon cancer tissues.

Moreover, the versatility of this method is noteworthy, as it transcends specific organ types. Its versatility stems from the fact that sample preparation, imaging, and network training do not involve organ-specific steps, allowing scalability to tissue slides from various organs. Similarly, our approach can extend to cytology slides as well, which also exhibit 3D structures but have conventionally been studied in a 2D context using BF microscopy.

Likewise, our framework demonstrates remarkable repeatability across different imaging devices. These results highlight the robustness of the proposed method, encompassing both holotomography and neural network architecture. This consistency emphasizes the potential for widespread adoption and utilization in diverse research and clinical settings.

While our proposed virtual staining method has been successfully applied to tissues with a thickness of up to 20 μm, the applicable thickness can be pushed further through technical advances of holotomography. Currently, holotomography faces challenges related to image degradation as the sample becomes thicker. The degradation in image quality can be attributed to the multiple scattering of the light within the thick sample, which hinders the accurate reconstruction of RI values through holotomography. Nonetheless, several strategies are being employed to address this challenge, including enhancements to optical conditions, the implementation of reconstruction algorithms tailored for thicker samples^32^, and the application of aberration correction algorithms^33^. With these strategies aimed at improving image quality in holotomography, we believe in a synergistic growth between virtual staining and holotomography.

Moreover, holotomography quantifies RI values, determining the protein concentration in specific regions. This empowers the proposed framework to conduct further quantitative analysis of subcellular structures. Specifically, segmentation or classification of cell types within the generated 3D H&E stained images can be performed using open-source, pre-trained resources^34^, with results directly applicable to the RI images for calculating 3D morphological and biophysical parameters. With these capabilities, we believe that the proposed framework can effectively overcome current limitations in histopathology, and broaden its utility across various histopathological studies, aiding in diagnosis and precision medicine.

## Methods

### Colon cancer slides preparation from Gangnam Severance Hospital

To create the training datasets, we utilized a 4 μm-thick colon cancer tissue slide prepared as tumor microarrays. The slide was newly diagnosed and reviewed by a pathologist (SJS) at Gangnam Severance Hospital in January 2023. Among 26 cases, a colon cancer tissue slide with grade two was randomly selected. From the chosen case, a formalin-fixed paraffin-embedded tissue block was obtained. The tissue block was sliced to a 4 μm thickness using a microtome. The slice was carefully placed on a slide glass and deparaffinized using xylene. Subsequently, the slide was then stained with H&E, mounted with a mounting medium, which has an RI value of 1.495 measured by a refractometer (R-5000, Atago), and finally covered with coverslips. Annotations of glands and stroma in the H&E stained images were carried out by an experienced pathologist (SJS), specializing in uropathology. This study was approved by the institutional review board (IRB) of Gangnam Severance Hospital (IRB No. 3-2022-0083) and conducted in accordance with the Declaration of Helsinki.

For the label-free thick tissue slides used in the testing phase, we randomly selected two cases of colon cancer. Formalin-fixed paraffin-embedded blocks from the selected cases were sliced into 10 and 20 μm-thick sections using a microtome, respectively. The subsequent procedures were consistent with those of training datasets, except that we did not stain the slides and mounted the slides immediately after the deparaffinization.

### Gastric cancer slides preparation from Mayo Clinic

For the training datasets, we obtained four gastric cancer tissue slides, each with a 5 μm thickness and H&E stain. The slide was newly diagnosed and reviewed by pathologists at the Mayo Clinic in January 2023. The slide preparation procedures were consistent with those conducted at Gangnam Severance Hospital. This study was approved by the institutional review board (IRB) of Mayo Clinic (IRB No. 3-2022-0083) and was conducted in accordance with the Declaration of Helsinki.

For testing purposes, a label-free, 20 μm-thick tissue slide was randomly selected from the gastric cancer cases. The slide was prepared using the same procedures as those performed at Gangnam Severance Hospital.

### Image acquisition

To measure the RI distributions of tissue slides, we employed a low-coherence holotomography setup (HT-X1, Tomocube). This system measures transmitted intensity images of a sample using four optimized Köhler illumination patterns^27^, from which the 3D RI distribution was reconstructed by deconvolution of the intensity images with theoretically estimated point spread functions. In the experimental setup, a light-emitting diode (LED) with a center wavelength of 449 nm within a digital micromirror device (DMD) module (DLP4500, Texas Instrument) was used as the illumination source. The blue wavelength was chosen to avoid overlap with the peak absorption spectrum of H&E staining. The illumination intensity was controlled using a DMD located in the Fourier plane, which was then projected onto the sample using a condenser lens (*f* = 180 mm, numerical aperture (NA) = 0.75). The diffracted light from the specimen was collected using an objective lens (numerical aperture (NA) = 0.95) and measured using a CMOS camera (FS-U3-28S5, FLIR) located in the image plane. The RI images offer a lateral and axial resolution of 155 nm and 1.07 μm, respectively. In addition to RI images, this system offers single-channel BF (scBF) images, which are single-intensity images taken under uniform circular pattern illumination. Wide-field RI images were obtained by stitching the tiled RI images of each field of view with the size of 227 × 227 μm. For the stitching, the raw intensity images were captured with an overlap of 23 μm for both RI and scBF images. To obtain the ground truth images, the H&E stained slides were scanned using the whole slide scanner manufactured by 3D HISTECH for colon cancer slides and by Leica for gastric cancer slides.

### All-in-focus

We applied an all-in-focus algorithm only to the RI images obtained from a slide used for network training^28^. This algorithm ensures that the single focal plane includes most of the cellular structures. As demonstrated in the previous study, we used a vignette size of 100 pixels with a step size of 10 pixels. Normalized variance was calculated in each window for all axial sections, and the window from the section with the maximum normalized variance value was selected as the best focal plane. Repeating the procedures for the whole 3D RI images resulted in a single all-in-focused RI image.

### Registration

Image registration between the RI and WSS images involved three steps. First, we manually cropped the regions from the WSS images where they were imaged using holotomography. Subsequently, we applied a spatial transform network^29^ to register the manually cropped WSS images with the scBF images acquired using holotomography. This step resulted in paired images of scBF and WSS at a wide-field scale. Then, we cropped the paired datasets into 1,024 × 1,024-pixel patches and repeated the registration using a spatial transform network between these patches to ensure precise alignment at each patch scale. Note that aligning the WSS images with scBF images directly leads to the alignment with RI images, as scBF and RI images are captured at the same position.

All the registration steps, except for the manual cropping, were conducted using spatial transform networks^29^. Specifically, EfficientNet^35^ was trained to discover the optimal affine transform matrix for the input WSS images, which will be used to transform the WSS images to be aligned with the target scBF images. The training was achieved by minimizing the loss defined using Pearson’s correlation coefficient (PCC):

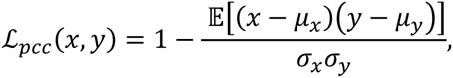

where *x* and *y* refer to the input WSS and target scBF images, respectively. *μ* and *σ* denote the mean and the standard deviation of each image, respectively. Once the training was completed, we employed the trained network to predict the optimized affine transform matrix, which was then applied to register the WSS images.

### Training details

A conditional GAN was used to train the network for generating 3D H&E images, primarily based on a previous study^36^. In this framework, the generator’s objective is to transfer the RI images into the feasible H&E stained BF images by minimizing the 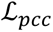 between the images generated by the network and their corresponding ground truth images. Simultaneously, the discriminator aims to distinguish the images from the network and genuine images when provided with the corresponding input images. The final objective of this framework becomes

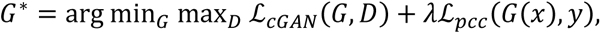

where 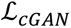 is defined as:

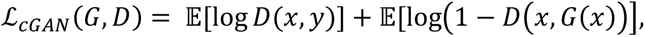

where *G*(·) and *D*(·) refer to the generator and discriminator network operators, respectively. *x* denotes an input image and *y* indicates a ground truth image.

For the architecture of the generator, we used SCNAS^30^, an optimized network specifically tailored for 3D medical image segmentation (Supplementary Fig. 2a). SCNAS employs a stochastic sampling algorithm within a gradient-based bi-level optimization framework to simultaneously search for the optimal network parameters at multiple levels using generic 3D medical imaging datasets. This search resulted in the discovery of a U-Net-like encoder-decoder structure with skip connections. In each micro-level architecture, we added a squeeze excitation block^37^. The key network parameters were as follows: activation function, leaky ReLU; normalization function, instance normalization; the size of the initial feature map, 12; the number of layers, 8; feature map multiplier, 2.

The discriminator consists of five convolutional layers with a kernel size of 4 and a stride of 2 (Supplementary Fig. 2b). The first four layers use leaky ReLU with a slope of 0.2, while the last layer employs ReLU as an activation function. Following the last layer, the result is activated by a Sigmoid function. The first and last layers use batch normalization, while the rest of the layers use instance normalization.

Both the generator and discriminator were trained using an adaptive moment estimation optimizer (ADAM) to update the learnable parameters^38^. Image augmentation techniques, such as adding blur, adjusting brightness, and introducing Gaussian noise, were randomly added to the training patches. A learning rate of 1 × 10^-4^ was maintained throughout the training process. Training was conducted for 40 epochs with a batch size of 1, and early stopping was employed to prevent overfitting. The network was implemented using Python version 3.8.0 and PyTorch^39^ version 1.13.1, and the procedures were executed on NVIDIA GeForce 4090 GPU.

The training results are compared with the ground truth images using SSIM, which is defined as below:

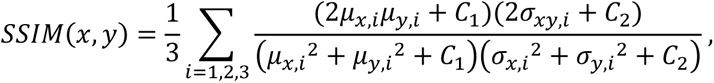

where *x* and *y* are the network-generated output and corresponding ground truth image, respectively. *μ* and *σ* refer to the mean and the standard deviation of each image, respectively, where an index *i* refers to the RGB channels of each image. σ_xy,i_ represents the covariance of both images calculated from *i*-th image channels. *C*_1_ and *C*_2_are regularization constants, set as 6.5025 and 58.5225, respectively, by default. The SSIM values were calculated for the selected windows. To make the spatial frequency range identical between the images, we performed low-pass filtering with the same kernel size for both images before calculating SSIM.

## Acknowledgement

This work was supported by the National Research Foundation of Korea (2015R1A3A2066550, 2022M3H4A1A02074314, RS-2023-00241278), an Institute of Information & Communications Technology Planning & Evaluation (IITP; 2021-0-00745) grant funded by the Korea government (MSIT), a KAIST Institute of Technology Value Creation, Industry Liaison Center (G-CORE Project) grant funded by MSIT (N10240002), the Korea Health Technology R&D Project through the Korea Health Industry Development Institute (KHIDI), funded by the Ministry of Health & Welfare, Korea (HI21C0977, HR22C1605).

---

**Supplementary Fig. 1.**
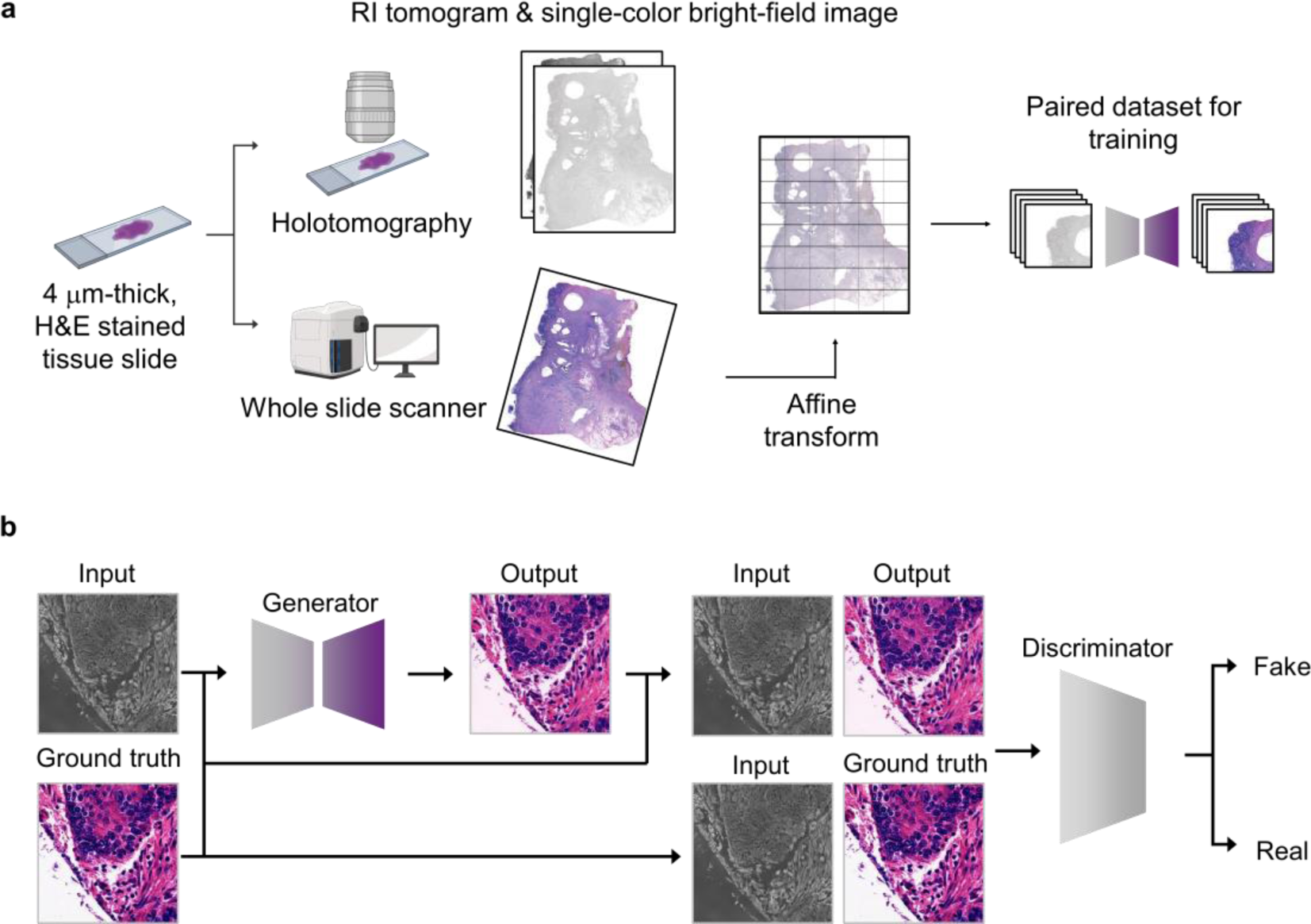
Dataset preparation and conditional generative adversarial network. **a,** Workflow of dataset preparation for the training phase. A conventional 4 μm-thick, H&E stained colon cancer tissue slides are imaged under holotomography and a whole slide scanner. Then, the images are registered using a spatial transform network^29^, resulting in a paired dataset for supervised learning. **b,** Overview of the conditional generative adversarial network^36^. The generator creates H&E stained images from input RI images, while the discriminator is trained to distinguish between the generated images and ground truth images.

**Supplementary Fig. 2.**
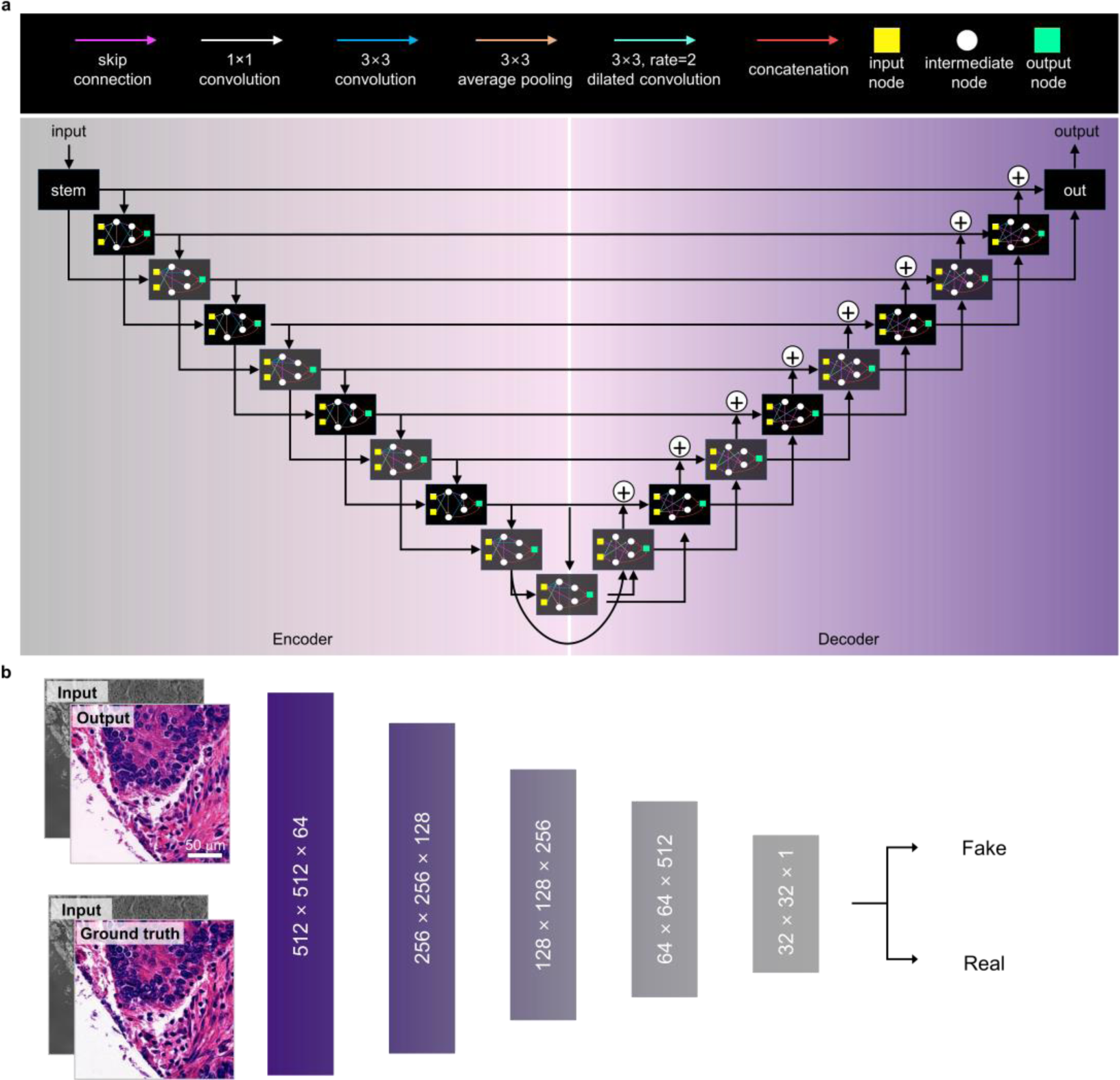
Network architecture. **a,** Architecture of the SCNAS used for generator^30^. The structure includes eight encoding micro-level architectures, a bridge, and another decoding eight micro-level architectures, resulting in a U-Net-like structure. **b,** Architecture of the discriminator. The structure is benchmarked from the previous work^36^, which includes five sequential convolutional layers.

**Supplementary Fig. 3.**
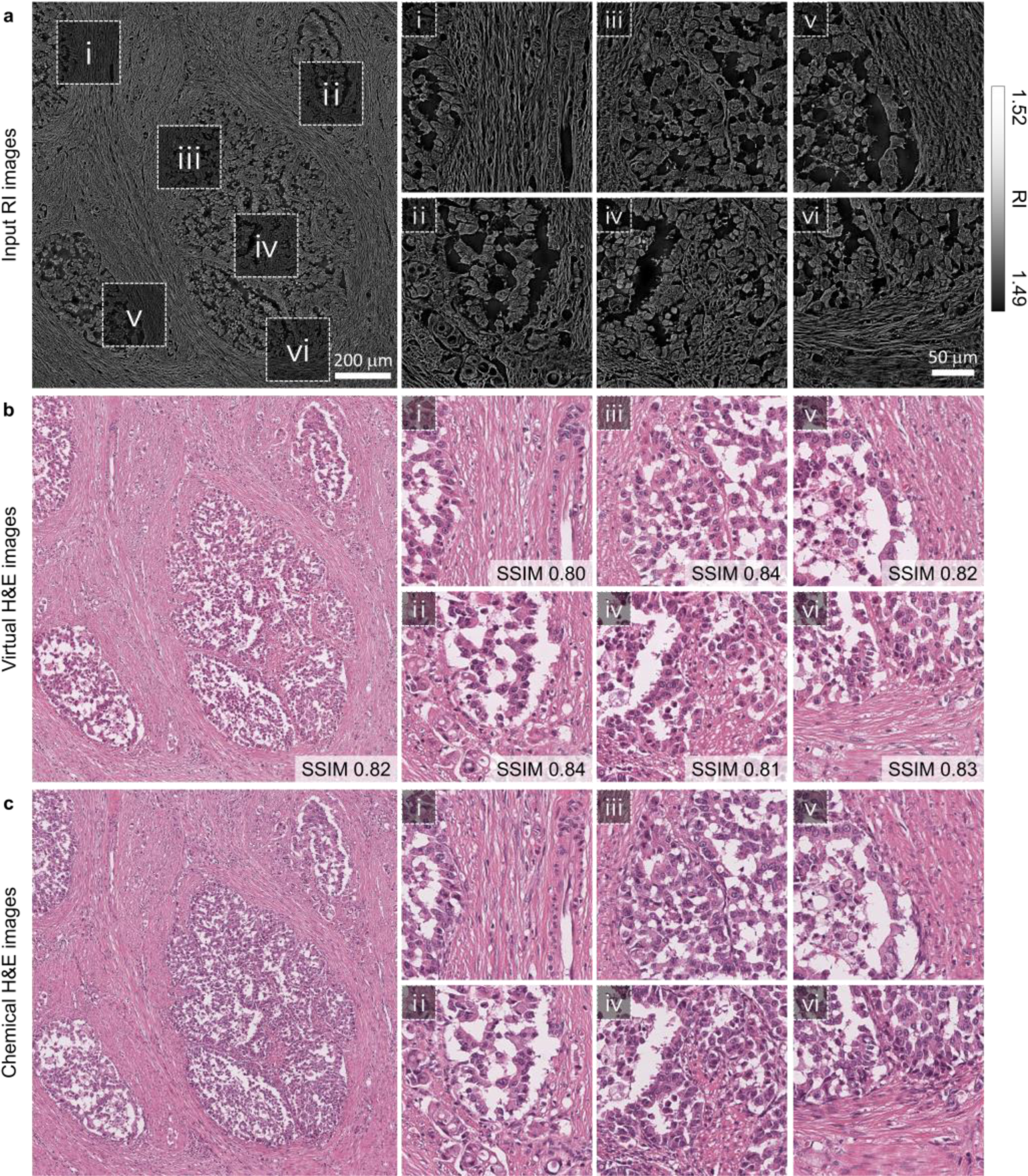
Validations of the trained network with a 5 μm-thick, H&E-stained gastric cancer tissue slide. **a,** Wide-field RI image obtained from a 5 μm-thick, H&E-stained gastric cancer slide and its detailed images. **b,** Wide-field H&E stained images generated by the trained neural network and its detailed images. **c,** Ground truth H&E images obtained using a WSS and its detailed images.

